# Comparative genomic analyses shed light on the introduction routes of rice-pathogenic *Burkholderia gladioli* strains into Bangladesh

**DOI:** 10.1101/2025.11.17.688954

**Authors:** Ismam Ahmed Protic, Md. Nasir Uddin, Andrew Gorzalski, Md. Rashidul Islam, David Alvarez-Ponce

## Abstract

**Background:** The gram-negative bacterium *Burkholderia gladioli* can cause diseases in a wide range of hosts, including humans and plants. In rice, it produces Bacterial Panicle Blight (BPB), an emerging disease that is threatening the production of this staple food worldwide. Recently, BPB has affected several rice fields in Bangladesh, and *B. gladioli* has been reported as the causative agent in one of these fields, raising questions about how the pathogen was introduced into the country. Understanding the introduction routes into Bangladesh and the pathogenicity mechanisms of *B*. *gladioli* is essential to develop sustainable management strategies to control BPB.

**Results:** To investigate the origin of pathogenic *B*. *gladioli* strains and their introduction routes into Bangladesh, we sequenced the complete genomes of 19 strains isolated from symptomatic rice panicles from four major rice-growing districts of the country (Rangpur, Faridpur, Natore, and Mymensingh). A phylogenetic analysis encompassing all 320 publicly available *B. gladioli* genomes along with the 19 newly sequenced genomes revealed five independent introduction events into Bangladesh—four of which showed close phylogenetic links to clinical isolates from the USA—and no evidence of inter-district dissemination within the country. Many host transitions (35 human-to-plant and 1 plant-to-human) were inferred across the phylogeny, highlighting the pathogen’s versatile and cross-kingdom transmission dynamics. Analysis of the presence/absence patterns of 37 virulence-related genes across all 339 *B. gladioli* strains indicated that human-to-plant host transitions often coincided with loss of the *tsr* and *motB* genes—which are associated with bacterial motility—, suggesting convergent adaptive shifts in response to host switches.

**Conclusion:** Our findings shed light on the introduction pathways of pathogenic *B*. *gladioli* strains into Bangladesh and highlight specific virulence-related genes as potentially responsible for cross-kingdom host shifts.

## INTRODUCTION

The genus *Burkholderia* is a group of gram-negative non-fermenting bacteria with more than 60 species that occur worldwide in virtually all possible environments, including the rhizosphere, plant roots, and water [1–4]. These include plant, animal, and human pathogenic bacteria, as well as environmentally beneficial species [5]. Some of these species can cause plant diseases, chronic lung infections in cystic fibrosis patients, and foodborne illnesses [6, 7]. The ecological versatility presented by many of the species may be due to their evolutionarily plastic and unusually large, multi-chromosome genomes [8, 9].

Some strains of *Burkholderia gladioli* act as plant pathogens, whereas others act as opportunistic human pathogens. *B. gladioli* was originally isolated as a pathogen of plants in the genus *Gladiolus* [10]. Four pathovars of *B*. *gladioli* with different host ranges have been reported so far. *B*. *gladioli* pv. *alliicola* causes onion bulb rot, *B*. *gladioli* pv. *gladioli* causes gladiolus rot, *B*. *gladioli* pv. *agaricicola* causes mushroom rot, and *B*. *gladioli* pv. *cocovenenans* causes serious food poisoning in humans [11–14]. *B*. *gladioli* not only causes significant agricultural losses worldwide, but also poses a threat to immunocompromised individuals, including cystic fibrosis patients and newborn infants [15, 16].

In rice, *B*. *gladioli* causes bacterial panicle blight (BPB) and poses a major threat to the rice industry in many countries [17]. *B. gladioli* was initially identified as a BPB pathogen in Japan and the Philippines in 1996, and later reported in the United States (Louisiana, 2001; Arkansas, 2009; Mississippi, 2012), Panama (2007), Ecuador (2014), China (2018), and Malaysia (2023) [18–26]. Two other species of *Burkholderia* can also cause BPB: *B*. *glumae* and *B*. *plantarii* [27, 28]. BPB is a seed-borne disease that causes several types of damage, including seedling blight, sheath rot, floret sterility, spotted grains, dark brown chaffy grains, and milling quality reduction, resulting in yield reductions of up to 75% [29–31]. It caused an estimated $61 million in losses in the Mid-South U.S. from 2003 to 2013 [32].

Bangladesh, the third-largest rice producer globally, is particularly vulnerable to BPB due to the significant contribution of rice to its national income. BPB was first reported in Bangladesh in 2023 from rice panicles of the local variety (Haridhan) from the Mymensingh district [31] and more recently from rice panicles of the Swarna variety from the Natore district in 2024 [33, 34], and the causal agents were determined to be *B*. *gladioli* and *B*. *glumae*, respectively. Interestingly, *B*. *gladioli* appears to be more prevalent than *B*. *glumae* in the country [31]. Among Bangladesh’s neighboring rice-producing countries—India, Myanmar, and Pakistan—only India has reported BPB, with *B*. *glumae* as the causal agent [35]. Therefore, the higher prevalence of *B*. *gladioli* over *B*. *glumae* in Bangladesh was unexpected. This raises critical questions regarding how *B*. *gladioli* was introduced into Bangladesh.

The variability in the virulence and host range of *B*. *gladioli* indicates the existence of intricate molecular processes that regulate host specificity. Although some virulence factors such as toxoflavin and type II, type III, and type VI secretion systems (T2SS, T3SS and T6SS) are highly conserved between *B*. *gladioli* and the well-studied rice pathogen *B*. *glumae* [36–38], the genetic determinants of host specificity in *B*. *gladioli* remain unclear. Gaining a deeper understanding of *B*. *gladioli*’s virulence mechanisms, prevalence and routes of introduction into Bangladesh is essential for developing effective strategies for controlling BPB and safeguarding global rice production. For instance, it could aid in strengthening the country’s quarantine system against potential entry points from neighboring regions.

In this study, we sequenced 19 pathogenic *B. gladioli* strains isolated from four major rice-growing districts in Bangladesh. To investigate the routes of introduction of the pathogen into the country, we performed phylogenetic analyses on 339 *B*. *gladioli* genomes (19 from our study plus 320 publicly available ones). We also investigated presence/absence and copy number variation (CNV) of 37 pathogenicity-related genes in order to understand the molecular determinants of strains’ host specificity (plants vs. humans).

## MATERIALS AND METHODS

### Sample collection and molecular identification

Rice fields cultivating eight different cultivars with typical BPB symptoms from 20 districts of Bangladesh were surveyed from mid-October 2022 to December 2023. A total of 300 fields (three locations from each district and five fields from each location) were sampled. From each field, panicles with typical BPB symptoms were collected. From each sample, we obtained isolates of the pathogen(s) causing the specific BPB symptoms following Islam et al. [22]. From each sample, 1g of symptomatic rice grains was surface sterilized, immersing it in 70% ethanol for 15 seconds, and then in 3% sodium hypochlorite for 1 min. Surface sterilized grains were ground in a mortar and 20 µl of extracted suspension was plated onto S-PG medium, which is specific for *B. gladioli* and its close relatives [29]. Purple-colored single colonies were streaked onto King’s B agar (KBA) [39] and were visually checked for the production of toxoflavin. Isolates producing a yellowish pigment (toxoflavin) on KBA medium were selected and purified as potential *B. gladioli* isolates. Genomic DNA was extracted using the Wizard® Genomic DNA Purification Kit (Promega, Madison, WI, USA). For molecular characterization, the 16S rDNA gene was amplified at around 1400 bp using the 16SF (5′-AGAGTTTGATCCTGGCTCAG-3′) and 16SR (5′-GGCTACCTTGTTACGACTT-3′) primers [17], and the *gyrB* gene was amplified at 479 bp using the *B. gladioli*-specific gla-FW (5’-CTGCGCCTGGTGGTGAAG–3’) and gla-RV (5’-CCGTCCCGCTGCGGAATA-3’) primers [40]. The isolates that were preliminary identified as *B. gladioli* according to these tests were used in rice pathogenicity tests both in field and net house conditions following refs. [17, 32], and Koch’s postulates were fulfilled by reisolating the same bacterium from the inoculated rice plants.

### Genome sequencing, assembly, and analysis

Forty six of our isolates tested positive for colony morphology, toxoflavin production, 16S amplification, and species-specific *gyrB* amplification and all of these isolates tested positive in pathogenicity tests. Their extracted DNA was subjected to quality assurance by Nanodrop (Thermo Fisher Scientific, USA), using the standard requirement of OD_260/280_ between 1.8 and 2.0, and OD_260/230_ between 2.0 and 2.2. Five of the isolates were discarded due to low DNA concentration prior to sequencing. Whole genome sequencing was conducted using paired-end reads (2×150bp) on the Illumina NextSeq 2000 platform at the Nevada Genomics Center. The raw sequence reads were imported into a Terra (https://app.terra.bio) workspace and analyzed with the TheiaProk_Illumina_PE_PHB v1.2.0 workflow under default settings [41]. The workflow performed quality control, read and adapter trimming, genome assembly, and taxon assignment of Illumina paired-end reads using fastq-scan [42], Trimmomatic [43], Shovill [44], and GAMBIT [45], respectively. Genome annotation was performed using Prokka v.1.14.5 [46]. The quality of sequenced genomes was further confirmed by BUSCO v.5.7.0 [47]. Nineteen of our 41 assemblies exhibited a BUSCO completeness above 90% and were unambiguously confirmed as *B. gladioli* by GAMBIT (Table 1).

**Table 1:** Summary genome sequencing statistics for 19 *B. gladioli* strains isolated from rice grains from Bangladesh.

| Isolate ID | Origin of isolate/district | Genome size (Mb) | N50 | Genome coverage | %GC | %BUSCO completeness |
| --- | --- | --- | --- | --- | --- | --- |
| BDBgla38A | Rangpur | 8.04 | 11993 | 21.4x | 67.91 | 97.00 |
| BDBgla41A | Rangpur | 8.32 | 47645 | 72.5x | 67.91 | 99.70 |
| BDBgla59A | Rangpur | 8.02 | 28445 | 35.2x | 67.84 | 99.30 |
| BDBgla78A | Faridpur | 8.16 | 60484 | 98.3x | 67.96 | 99.60 |
| BDBgla79A | Faridpur | 8.07 | 20227 | 40.7x | 67.88 | 98.40 |
| BDBgla81A | Faridpur | 8.11 | 36331 | 40.5x | 67.95 | 99.70 |
| BDBgla83A | Faridpur | 7.73 | 7913 | 20.0x | 67.98 | 92.00 |
| BDBgla117A | Natore | 8.38 | 38907 | 67.7x | 67.88 | 99.60 |
| BDBgla119A | Mymensingh | 8.43 | 40496 | 69.1x | 67.76 | 99.30 |
| BDBgla120A | Mymensingh | 8.42 | 37957 | 65.9x | 67.76 | 99.70 |
| BDBgla122A | Mymensingh | 8.40 | 34679 | 55.8x | 67.77 | 99.00 |
| BDBgla130A | Mymensingh | 8.40 | 26019 | 54.2x | 67.73 | 99.10 |
| BDBgla132A | Mymensingh | 8.38 | 32382 | 42.8x | 67.78 | 99.10 |
| BDBgla135A | Mymensingh | 8.47 | 46671 | 44.2x | 67.73 | 99.60 |
| BDBgla136A | Mymensingh | 8.82 | 56072 | 59.0x | 67.43 | 100.00 |
| BDBgla140A | Mymensingh | 8.44 | 63996 | 66.4x | 67.79 | 100.00 |
| BDBgla150A | Mymensingh | 8.43 | 57006 | 64.5x | 67.78 | 99.70 |
| BDBgla151A | Mymensingh | 8.42 | 57209 | 59.4x | 67.79 | 99.70 |
| BDBgla162A | Mymensingh | 8.40 | 49226 | 43.7x | 67.79 | 99.60 |
| Average |  | 8.31 | 39666.21 | 53.8x | 67.81 | 98.95 |

We obtained all 331 *B. gladioli* genome assemblies available in GenBank [48]. Eleven of these assemblies were discarded due to anomalies in genome length and/or contamination during sequencing. In addition, strain SMAGNO155 was excluded since it introduced a large branch into our phylogenetic tree; however, the exclusion of SMAGNO155 did not affect the topology of the tree. All the retained assemblies from GenBank had an N50 ≥ 13,124. We assembled the genome of strain BCC1317 (the only sequenced strain from an industrial setting) from its raw sequence data and added it to our analysis. The resulting dataset thus contained 339 *B. gladioli* genomes (19 from our study, 319 from GenBank, and 1 assembled from SRA data; Table S1).

For each pair of genomes, we calculated the average nucleotide identity (ANI) using FastANI v1.34 [49]. A heatmap was generated using ANIclustermap v1.4 [50]. The core genome present in all *B*. *gladioli* strains was inferred using Roary v3.6.0 [51]. FastTree v2.0.0 [52] was used to build a maximum-likelihood phylogenetic tree from the core gene alignment using the generalized time-reversible (GTR) model and 1000 bootstraps. To root our tree, we conducted a separate phylogenetic analysis using the same methods, including *B*. *glumae* BGR1 as an outgroup. The resulting tree was visualized using Interactive Tree of Life (iTOL) v6 [53].

### Identification of pathogenicity genes

We obtained the proteomes from the 339 *B. gladioli* strains included in our analysis from different sources. A total of 313 were obtained from the RefSeq database [54], six were obtained from GenBank (strains KRS027, SOL3_8, SOL2_4, SH2_2, SH3_3, and SH1_3), and for the remaining 20 (strain BCC1317 plus the 19 strains from Bangladesh) we used our Prokka annotations. The proteome of *B. glumae* BGR1 was also obtained from the RefSeq database.

For each *B. gladioli* proteome, we identified orthologs by running OrthoMCL v.2.0.9 [55] against the *B*. *glumae* BGR1 proteome with E-value < 10^-5^ and match length ≥ 50%. We used *B. glumae* BGR1 as reference because plant pathogenic *B. gladioli* strains are not well characterized, the two species are closely related [56], and BGR1 is the most studied *Burkholderia* plant pathogen and the most studied *B. glumae* strain [9, 36, 37].

After genome-wide orthologs identification, we focused our analyses on 37 specific genes that are important for plant pathogenesis: those responsible for producing toxoflavin (13 genes distributed in 2 clusters in BGR1 [57]), lipase (2 genes located in tandem [58]) and polygalacturonase (2 genes located in separate chromosomes [59]), and 20 genes involved in motility (10 genes involved in flagella production, 8 genes involved in chemotaxis, and 2 genes involved in other motility processes, all distributed in 2 clusters [60–65]) (Table 2).

**Table 2:**
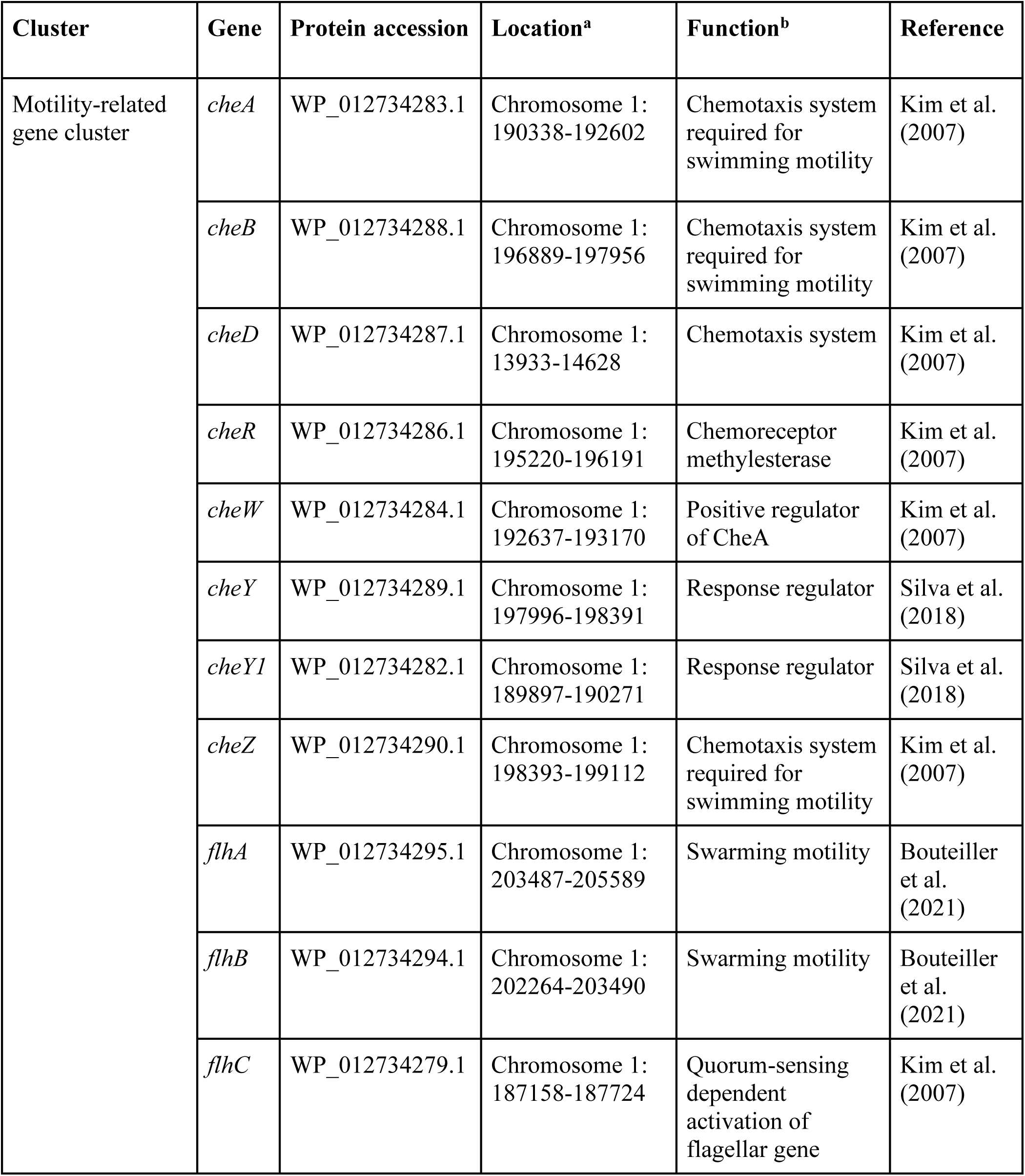
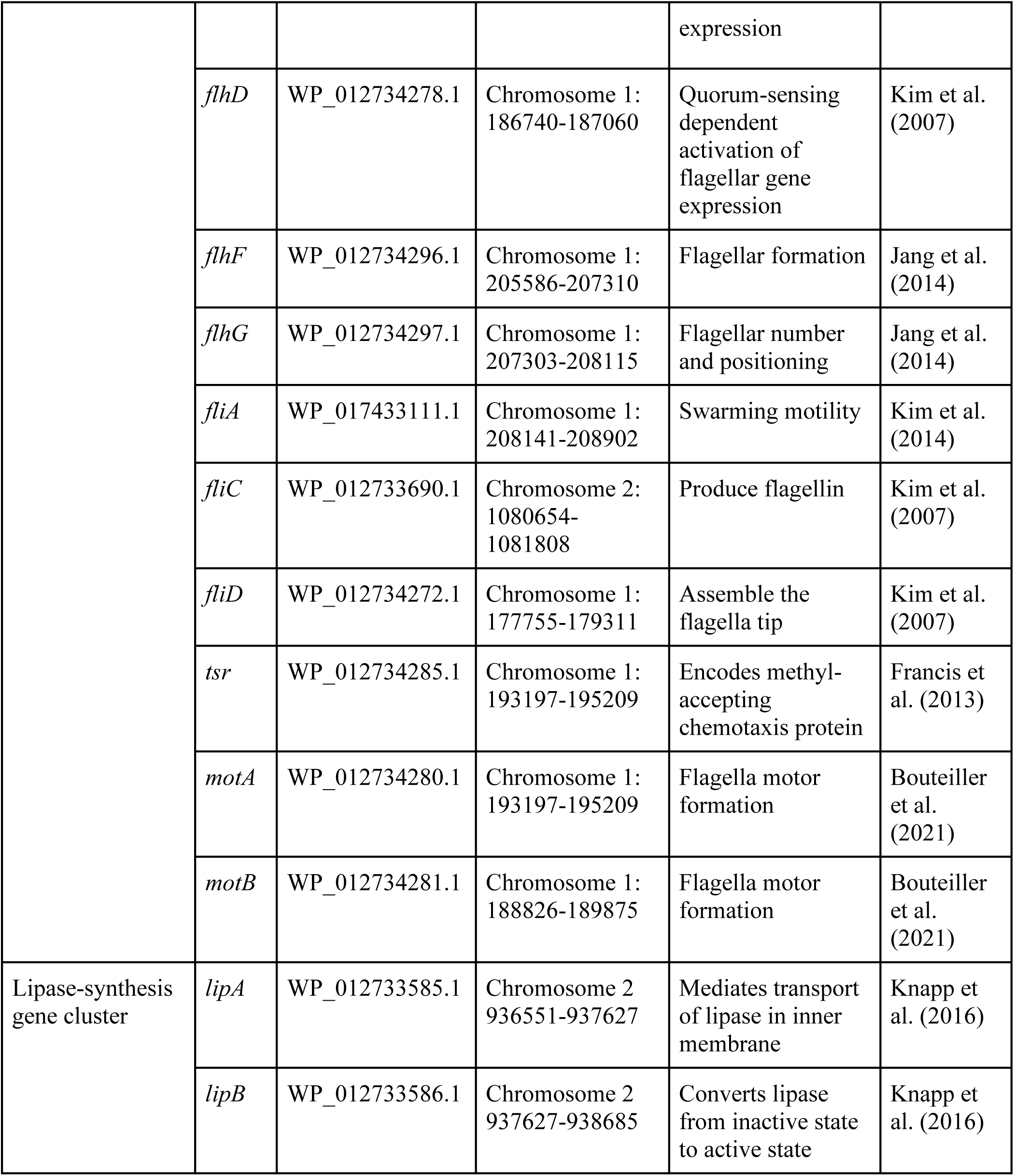
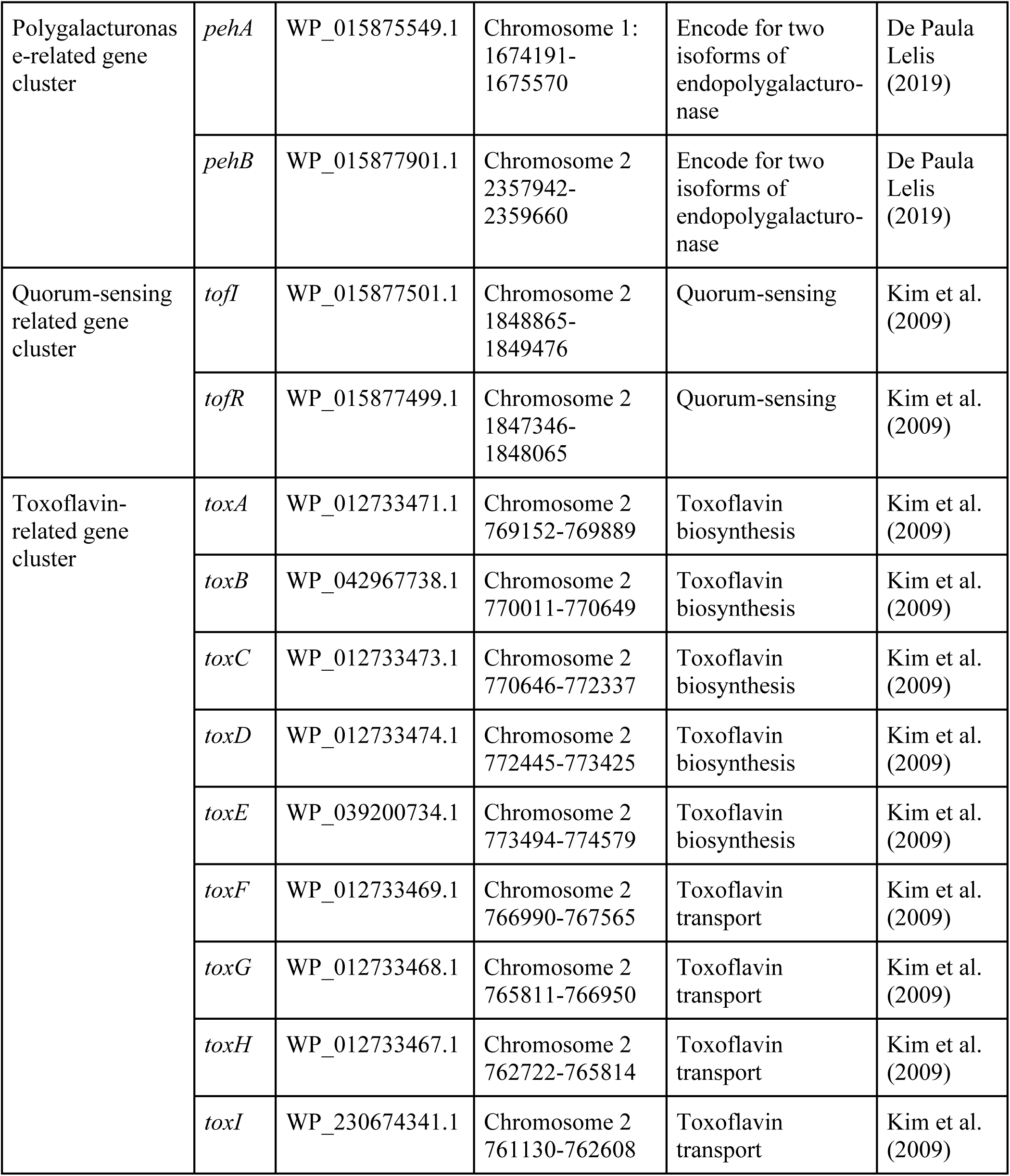
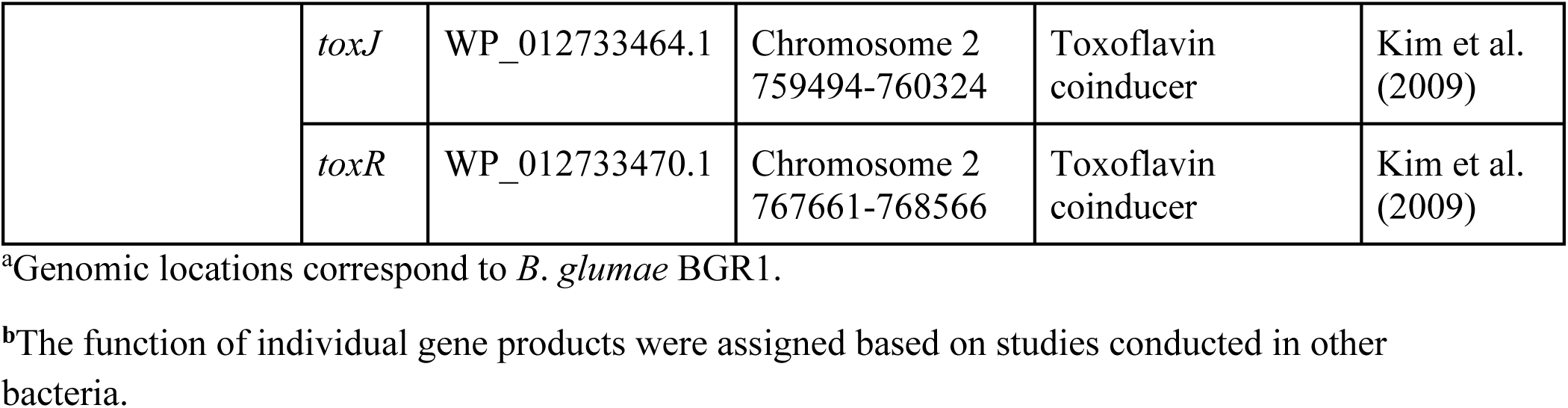
Virulence-related genes included in our study.

### Evaluation of the association between gene gains/losses and host switches

For each of the 37 genes associated with plant and/or human pathogenicity (Table 2), we used the Fisher’s exact test (FET) to test whether the fraction of strains containing the gene was significantly different among plant- and human-derived strains, finding significant differences in five genes (*pehB*, *toxI*, *tsr*, *cheR*, and *motB*, *P* < 0.05; Table 3). We then tested whether human-to-plant host switches tended to occur in the same branches of the phylogeny as changes in presence/absence (gains/losses) of these five genes. To that end, we trimmed our tree to include only plant- and human-derived *B. gladioli* strains (n = 322). We then used maximum parsimony to infer the branches in which there were host changes (we identified 35 human-to-plant transitions and one plant-to-human transition) or changes in presence/absence of the 5 genes. For each gene, we used as statistic *X*—the number of branches in which both the host switched from human to plant and the gene switched from absent to present (if the gene was more frequent among plant-derived strains) or from present to absent (if the gene was more frequent among human-derived strains). We then used a Monte Carlo method to estimate the null distribution of *X*. For each gene, we conducted 10,000 simulations. In each simulation, a number of presence→absence or absence→presence changes (equal to the observed number of changes) were randomly assigned to branches. The likelihood of a gene changing state in any branch was proportional to the branch’s length. For each gene, we computed a *P*-value as the fraction of simulations in which *X* was higher or equal to the observed *X* value.

**Table 3:** Comparison of gene presence between human and plant-originated *B*. *gladioli* strains.

| **Gene** | **% Present in human-originated strains** | **% Present in plant-originated strains** | ***P*-value<sup>a</sup>** | ***Q*-value<sup>b</sup>** |
| --- | --- | --- | --- | --- |
| *cheA* | 100.00 | 100.00 | 1.00 | 1.00 |
| *cheB* | 99.20 | 100.00 | 1.00 | 1.00 |
| *cheD* | 99.20 | 98.63 | 0.54 | 1.00 |
| *cheR* | 99.60 | 87.67 | 8.41×10<sup>-6</sup>* | 7.78×10<sup>-5</sup>* |
| *cheW* | 100.00 | 100.00 | 1.00 | 1.00 |
| *cheY* | 100.00 | 100.00 | 1.00 | 1.00 |
| *cheYI* | 98.40 | 97.26 | 0.62 | 1.00 |
| *cheZ* | 100.00 | 100.00 | 1.00 | 1.00 |
| *flhA* | 99.60 | 100.00 | 1.00 | 1.00 |
| *flhB* | 99.60 | 100.00 | 1.00 | 1.00 |
| *flhC* | 100.00 | 100.00 | 1.00 | 1.00 |
| *flhD* | 100.00 | 100.00 | 1.00 | 1.00 |
| *flhF* | 99.20 | 100.00 | 1.00 | 1.00 |
| *flhG* | 100.00 | 100.00 | 1.00 | 1.00 |
| *fliA* | 100.00 | 100.00 | 1.00 | 1.00 |
| *fliC* | 100.00 | 97.26 | 0.05 | 0.31 |
| *fliD* | 100.00 | 98.63 | 0.23 | 1.00 |
| *tsr* | 100.00 | 73.97 | 7.02×10<sup>-14</sup>* | 2.60×10<sup>-12</sup>* |
| *motA* | 100.00 | 100.00 | 1.00 | 1.00 |
| *motB* | 100.00 | 76.71 | 2.13×10<sup>-12</sup>* | 3.94×10<sup>-11</sup>* |
| *lipA* | 100.00 | 100.00 | 1.00 | 1.00 |
| *lipB* | 100.00 | 98.63 | 0.23 | 1.00 |
| *pehA* | 85.60 | 83.56 | 0.71 | 1.00 |
| *pehB* | 4.00 | 23.28 | $2.63 \times 10^{-6}$* | $3.25 \times 10^{-5}$* |
| *tofI* | 98.80 | 97.26 | 0.32 | 1.00 |
| *tofR* | 94.00 | 91.78 | 0.58 | 1.00 |
| *toxA* | 99.20 | 100.00 | 1.00 | 1.00 |
| *toxB* | 99.20 | 100.00 | 1.00 | 1.00 |
| *toxC* | 98.00 | 100.00 | 0.59 | 1.00 |
| *toxD* | 99.60 | 100.00 | 1.00 | 1.00 |
| *toxE* | 100.00 | 100.00 | 1.00 | 1.00 |
| *toxF* | 98.80 | 100.00 | 1.00 | 1.00 |
| *toxG* | 98.80 | 100.00 | 1.00 | 1.00 |
| *toxH* | 99.60 | 100.00 | 1.00 | 1.00 |
| *toxI* | 100.00 | 94.52 | $2.00 \times 10^{-3}$* | $1.81 \times 10^{-2}$* |
| *toxJ* | 100.00 | 100.00 | 1.00 | 1.00 |
| *toxR* | 99.20 | 100.00 | 1.00 | 1.00 |

<sup>a</sup>Calculated using the Fisher's exact test.
<sup>b</sup>False Discovery Rate calculated using the Benjamini-Hochberg method.
\*, Statistically significant ( $P < 0.05$ or $Q < 0.05$ ).

## RESULTS

### Molecular identification, genome sequencing, and taxonomy of *B*. *gladioli* isolates

We sampled rice panicles displaying typical BPB symptoms from 300 rice fields from 20 rice-growing districts in Bangladesh. From each sample, we obtained a bacterial isolate. Out of the 300 isolates, 46 were preliminarily identified as *B. gladioli* based on their characteristic colony morphology on S-PG medium, production of the yellow pigment toxoflavin on King’s B agar, and 16S and *gyrB* PCR amplification. The remaining 254 isolates failed at least one of these tests, indicating that they belong to non-*Burkholderia* or to other *Burkholderia* species. All 46 isolates preliminarily identified as *B. gladioli* tested positive in pathogenicity tests in field and net house conditions. Five of these isolates were discarded due to low DNA concentration prior to sequencing. In total, 41 isolates were sequenced, and 19 of them (3 from the Rangpur district, 4 from the Faridpur district, 1 from the Natore district, and 11 from the Mymensingh district) were confirmed as *B. gladioli* based on taxonomy prediction by GAMBIT, and genome assembly completeness from BUSCO reports (cutoff: >90% completeness). The remaining 22 samples were classified as unclassified members of the *Burkholderia* genus (*n* = 14) or as members of other genera (*n* = 8). The sizes of the 19 *B. gladioli* genomes ranged from 7.74 Mb (BDBgla83A) to 8.82 Mb (BDBgla136A), with an average of 8.31 Mb, which falls within the previously reported genome size range for *B*. *gladioli* strains (7.32–8.94 Mbp [2]), and the sequencing depth ranged from 20.0× (BDBgla83A) to 98.3× (BDBgla78A), with an average of 53.8× (Table 1).

To provide a comprehensive understanding of the genetic variability of *B*. *gladioli*, a total of 320 publicly available genomes were also analyzed. The majority of the isolates (*n* = 249) were from clinical samples (from patients with cystic fibrosis), including 217 from the USA, 8 from the UK, 11 from Canada, 1 from Australia, and 1 from Italy. Fifty four of the isolates were derived from plants (27 from onion, 9 from rice, 7 from gladiolus, 5 from maize, 1 from Chapalote corn, 1 from fermented cornmeal, 1 from sugarcane, 1 from wheat, 1 from *Dendrobium officinale*, and 1 from oil palm); these isolates included 4 classified as *B. gladioli* pathovar *gladioli* and 1 classified as *B*. *gladioli* pv. *alliicola*. The remaining 17 isolates were derived from various sources (5 from environmental samples, 4 from soils, 3 from unknown sources, 2 from *Lagria villosa* beetles, 1 from an industrial site, 1 from a mosquito, and 1 from water) (Table S1).

All 339 genomes were confirmed as members of a single bacterial species by comparing the calculated ANI values (ranging between 97.5% and 99.93%) with the standard cut off (95%). Our ANI heatmap resulted in three distinct groups with >98% identity (Fig. 1). Since the results of our clustering analysis were consistent with those of a previous study of *B. gladioli* strains [5], we followed the same group nomenclature. Group 1 (*n* = 63) consisted of three closely related sub-groups, 1A (*n* = 23), 1B (*n* = 14) and 1C (*n* = 27). Groups 2 and 3 consisted of 116 and 159 strains, respectively. Strains isolated from plants distributed across all groups and subgroups: 6 in subgroup 1A (5 from maize and 1 from fermented cornmeal), 9 in subgroup 1B (8 from rice and 1 from oil palm), 17 in subgroup 1C (all 17 from rice), 22 in group 2 (19 from onion, 2 from rice, and 1 from gladiolus), and 19 in group 3 (8 from onion, 6 from gladiolus, 1 from Chapalote corn, 1 from *Dendrobium officinale*, 1 from rice, 1 from sugarcane, and 1 from wheat). Our 19 Bangladeshi strains clustered within subgroups 1B (*n* = 4) and 1C (*n* = 15).

**Figure 1:**
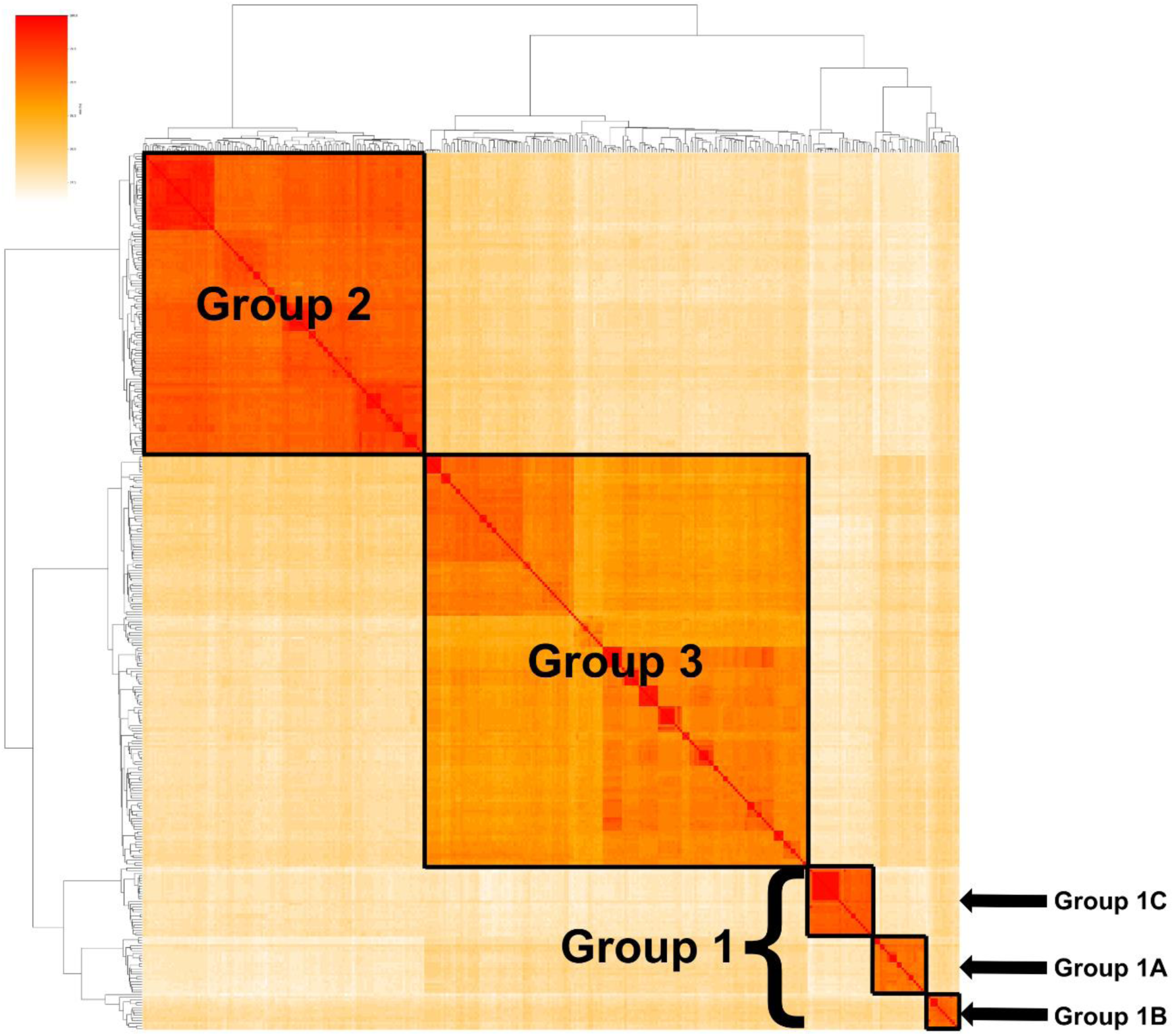
Heat map based on the ANI of 339 B. gladioli genomes. Darker shading indicates a greater percentage of identities. The black outlined boxes show sub-groups with greater than 97.3 % ANI . The main clusters are labeled as group 1, 2 and 3, with group 1 isolates subdividing further into subgroups 1A, 1B, and 1C (bottom right).

### Phylogenomic analysis

From the 339 *B. gladioli* genomes, we identified 3622 core genes (present in all of the strains). Using these genes, we obtained a phylogenetic tree to establish the phylogenetic position of our 19 Bangladeshi strains (Fig. 2). A separate tree including *B. glumae* BGR1 as an outgroup was generated to root our tree. The clusters defined based on the ANI values (groups 2 and 3 and subgroups 1A, 1B and 1C) corresponded to distinct clades in the tree. However, our phylogenetic analysis indicates that group 1 is polyphyletic. The most studied *B*. *gladioli* strain, BSR3, which causes BPB in rice, clustered in subgroup 1B.

**Figure 2:**
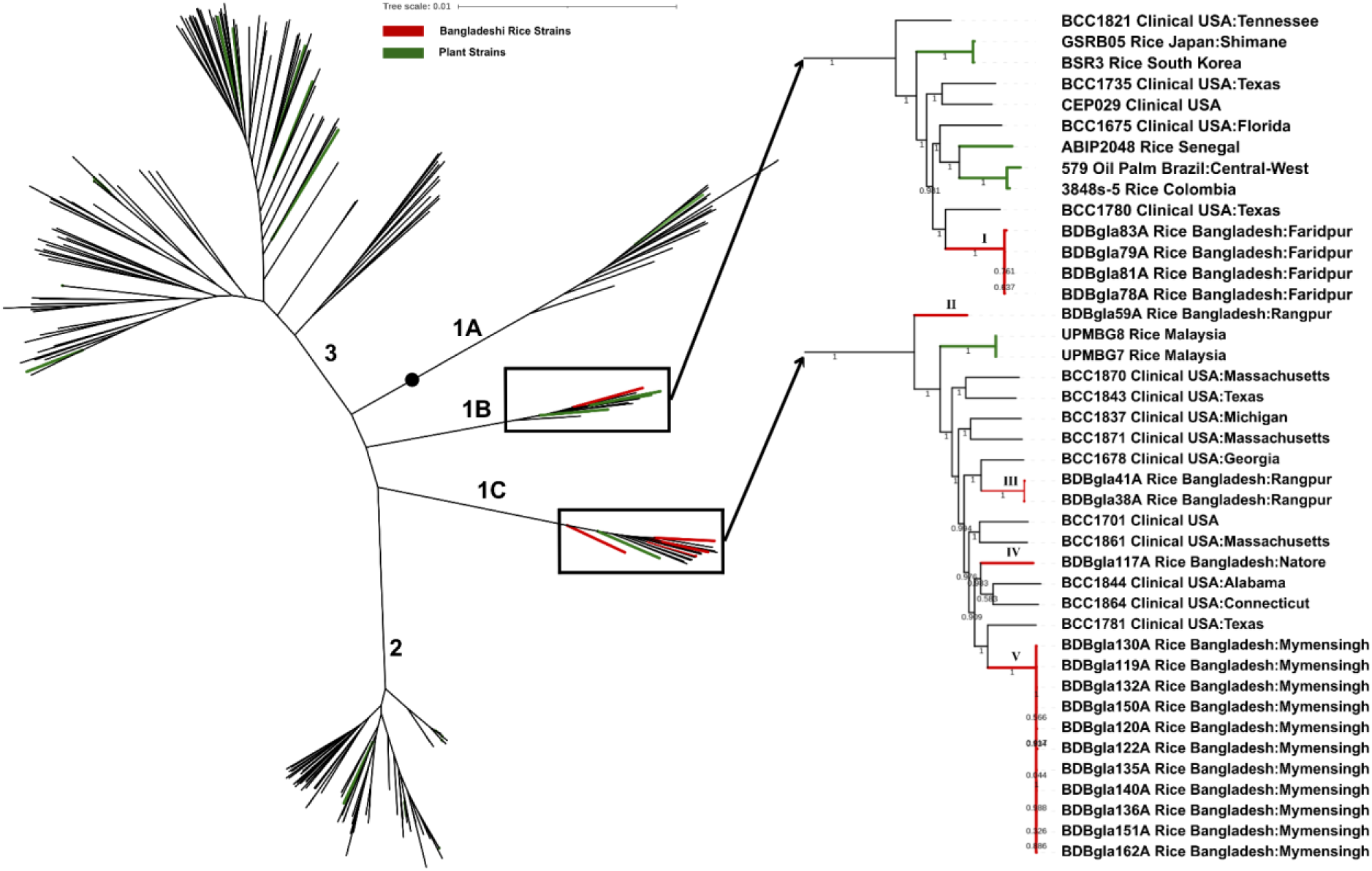
Maximum-likelihood phylogenetic tree based on the 3622 core genes of 339 *B*. *gladioli* strains. Green colored branches represent strains isolated from plants, and red colored branches represent strains isolated from rice displaying symptoms of BPB from Bangladesh. The five distinct Bangladeshi clades are labeled using Roman numbers (I-V). The tree was rooted using *B*. *glumae* BGR1 as an outgroup. The position of the root is represented by a black circle. The scale bar corresponds to 0.01 substitutions per site.

Our 19 Bangladeshi strains formed 5 different clades (I–V), suggestingthat pathogenic *B. gladioli* strains were introduced into Bangladesh at least 5 times independently. Each clade contained strains exclusively from a single district, indicating no inter-district spreading events within Bangladesh. Within subgroup 1B, BDBgla78A, 79A, 81A, and 83A (all from the Faridpur district) formed clade I, which had BCC1780 (a clinical sample from Texas, USA) as sister taxon. The clade including these 5 strains and BCC1780 was sister group to a clade including BCC1675 (also a clinical sample from Florida, USA) and three plant strains (ABIP2048, 3848s-5 and 579, isolated from rice from Senegal, rice from Colombia, and oil palm from Brazil, respectively). Subgroup 1B also included another two strains isolated from rice (GSRB05 and BSR3, isolated from Shimane, Japan, and from South Korea, respectively) and three clinical samples from USA patients (BCC1821, BCC1735, and CEP029).

Subgroup 1C contains four clades (II–V) of Bangladeshi strains. Clade II is composed of Bangladeshi strain BDBgla59A (from the Rangpur district), which was the sister taxon to all the other strains in subgroup 1C. The other three Bangladeshi clades (III–V) were embedded within a clade composed of clinical samples from the USA. First, Bangladeshi strains BDBgla38A and 41A (clade III; both from Rangpur) were the sister group of clinical sample BCC1678 from Georgia, USA. Second, Bangladeshi strain BDBgla117A (clade IV; from Natore) was sister taxon to a clade including clinical samples BCC1844 and BCC1864 (from Alabama and Connecticut, respectively). Third, Bangladeshi strains BDBgla119A, 120A, 122A, 130A, 132A, 135A, 136A, 140A, 150A, 151A and 162A (clade V; all from Mymensingh) were the sister group to clinical sample BCC1781 (from Texas). Subgroup 1C also included two strains isolated from rice in Malaysia (UPMBG8 and UPMBG7).

### Association of genes with hosts

For each *B. gladioli* proteome, we conducted a separate OrthoMCL analysis against the *B. glumae* BGR1 proteome. We then retained the results for 37 genes known to be involved in plant pathogenicity (Table 2). Nine of these genes were present in all 339 *B. gladioli* genomes, whereas the remaining 28 were absent from some of the genomes (Table S2).

For each gene, we compared the fraction of human-derived strains vs. the fraction of plant-derived strains in which the gene was present. Four of the genes (*toxI*, *tsr*, *cheR*, and *motB*) were significantly more represented among human-derived strains (100%, 100%, 99.60%, and 100%, respectively) than among plant-derived strains (94.52, 73.97%, 87.67%, and 76.71%, respectively; FET, *P* < 0.05). In contrast, *pehB* was more common among plant-derived strains than among human-derived strains (23.28% vs. 4%; *P* = 2.63×10^-6^; Table 3). These differences remain significant after controlling for multiple testing using the Benjamini-Hochberg approach [66] (Table 3).

However, the FET assumes that all observations are independent, not taking into account the phylogenetic relationships among the strains. This is problematic since closely related strains tend to exhibit similar genomes and a similar host specificity (e.g., ref. [67]). In addition, it does not take into account the number of switches in host specificity and in gene presence/absence. To address these limitations, we employed maximum parsimony to infer the minimum number of host shifts (*n* = 36, including 35 human-to-plant and 1 plant-to-human transitions) and the minimum number of changes in presence/absence of each of the five genes across the phylogeny (1 for *pehB*, 6 for *tsr*, 6 for *motB*, 8 for *cheR* and 2 for *toxI*). Next, for each gene, we computed *X*, the number of branches in which both the host switched from human to plant and the gene switched from absent to present (if the gene was more frequent among plant-derived strains) or from present to absent (if the gene was more frequent among human-derived strains), and used a Monte Carlo method to determine *X*’s statistical significance. The test was significant for two of the genes (*tsr* and *motB*, *P* = 0.0011), indicating that human-to-plant host transitions co-occurred with changes in presence/absence of these genes more often than expected based on a random evolution model (Table S3). The strains that lost *tsr* also lost *motB*, except for strains BCC1317, BDBgla140A and BDBgla150A, which lost *tsr* but not *motB*.

Out of the 37 genes analyzed, 31 exhibited no CNV (all genomes presenting the gene had the same number of copies), 3 exhibited subtle variation (less than 10% of the genomes presenting the gene had a number of copies that was different from the most common number of copies), and 3 exhibited substantial variation (more than 10% of the genomes presenting the gene had a number of copies that was different from the most common number of copies). Interestingly, most plant-derived strains retained *motB* and *tsr*, but these genes were absent from most Bangladeshi strains. Conversely, the *pehB* gene was present in most Bangladeshi strains but absent from most other plant-derived strains. None of the 37 genes analyzed exhibited CNV among the 19 *B. gladioli* strains from Bangladesh.

## DISCUSSION

Our study provides information about the origin of pathogenic *B*. *gladioli* strains and their likely introduction routes into Bangladesh, and helps understand the mechanisms of genomic adaptation of the pathogen to its plant hosts. We confirmed the presence of pathogenic *B*. *gladioli* in 19 fields from four major rice-growing districts in Bangladesh. Our phylogenomic analysis revealed that these strains clustered into five distinct clades, suggesting at least five independent introduction events into the country rather than a single introduction followed by a local expansion. The phylogenetic distribution of strains from clinical samples, plants, and the environment (plant-derived strains are scattered throughout the phylogeny), along with the numerous host shifts identified (35 human-to-plant and 1 plant-to-human transitions), highlights the pathogen’s versatile host associations and cross-kingdom adaptive dynamics. Further analysis of pathogenicity-related genes identified key associations between gene presence and host specificity.

The clear phylogenetic proximity of most Bangladeshi *B*. *gladioli* strains (4 out of 5 clades: I, III, IV, and V) to clinical isolates from the USA suggests that the introduction of the pathogen to Bangladesh is the result of international trade of agricultural products and/or other human-mediated transport from the USA on at least four occasions. For instance, clade I was sister group to the clinical sample BCC1780 from Texas, USA. Additionally, these strains, along with BCC1780, were sister group to a separate clade that includes BCC1675 (a clinical isolate from Florida, USA) and three plant strains—ABIP2048, 579, and 3848s-5—from Senegal, Brazil, and Colombia, respectively. Similarly, Bangladeshi strains BDBgla38A and 41A (clade III; both from Rangpur) formed a sister group to the clinical isolate BCC1678 from Georgia, USA. BDBgla117A (clade IV; from Natore) was a sister taxon to a clade comprising clinical isolates BCC1844 and BCC1864 from Alabama and Connecticut, respectively. Additionally, BDBgla119A, 120A, 122A, 130A, 132A, 135A, 136A, 140A, 150A, 151A, and 162A (clade V; all from Mymensingh) formed a sister group to the clinical isolate BCC1781 from Texas, USA. However, the putative origin of clade II is unclear based on our phylogenomic analyses, since it has as sister group a clade containing samples from different countries. The four potential introductions from the USA to Bangladesh, however, could have occurred through one or more intermediary countries rather than directly from the USA.

Notably, each Bangladeshi clade contained strains exclusively from a single district, indicating no inter-district spread within Bangladesh and supporting the idea of multiple, independent introduction events into the country rather than internal dissemination. Clades II and III were composed exclusively from strains from Rangpur, whereas clades I, IV and V were composed of strains from Faridpur, Natore and Mymensingh, respectively. These observations indicate that the pathogen was introduced at least once into Faridpur, once into Natore, once into Mymensingh, and twice into Rangpur independently. However, our current dataset includes only 19 genomes from four different districts, which limits our ability to draw definite conclusions. Additional sampling from other rice-growing regions across Bangladesh will be essential to obtain a more comprehensive understanding of genetic diversity, local evolution, and potential inter-district transmission dynamics of *B*. *gladioli*.

To further investigate the possible sources of *B. gladioli* introductions into Bangladesh, we examined Bangladesh’s rice import history, given that BPB is a seed-borne disease. Over the past decade, Bangladesh has been importing rice from India, Thailand, and Myanmar due to low tariffs and freight costs. According to the Grain and Feed report of the USDA, 71% of rice imports were from India, 17% from Thailand, and 12% from Myanmar during the 2022/23 marketing year [68]. This trend contrasts with previous years in which Bangladesh imported rice from other rice-producing countries like South Korea and Japan. For instance, in 2012, 2013, and 2015, rice imports from South Korea were valued at approximately US$20.9K, $22.3K, and $29.7K, respectively. Additionally, Bangladesh imported rice worth $1.4 million and $1.8 million from Japan in 2007 and 2008, respectively. The fastest-growing rice import markets for Bangladesh between 2022 and 2023 were Thailand ($14.3M), the United States ($21.2K), and Brazil ($1.8K) [69]. Although the Bangladeshi *B*. *gladioli* strains are phylogenetically linked to clinical isolates from the USA, before the recent trade history, there were no known rice imports between the USA and Bangladesh. Nonetheless, BPB was originally identified in Japan in 1956 [27]. Since then, it has been found in China, India, Thailand, and the United States, among other rice-producing nations [17, 25, 29, 35]. This illustrates how global trade may have contributed to the spread of *B*. *gladioli*.

The existence of closely related clinical, plant, and environmental isolates (Fig. 2) reinforces the idea that *B*. *gladioli* can transition among hosts. The survival of *Burkholderia cepacia* complex (Bcc) species for long periods of time on environmental surfaces, especially in sputum, suggests the possibility of indirect transmission. Indeed, natural environments provide potential reservoirs for the acquisition of the pathogen [70], and the transmission of *Burkholderia* among cystic fibrosis patients can occur through environments [71, 72]. In addition, rhizosphere, soil, and terrestrial freshwater environments are common sources of *Burkholderia* [73]. In hospital settings, *B. cepacia* has also been transmitted via contaminated tap water or patient materials disposed of in drains, highlighting the drainage system as a potential transmission pathway [74, 75]. Beyond clinical and environmental settings, a case study reported that the wild mushroom *Suillus luteus* can work as a potential reservoir of *B*. *gladioli*, and that contamination may occur directly or indirectly via animals. The study also confirmed that the strains isolated from *S. luteus* are invasive pathogens of fleshy fruits, vegetables, and edible mushrooms [76]. Furthermore, symbiotic associations with *B*. *gladioli* have been identified in at least six beetle species of the subfamily Lagriinae (Tenebrionidae), in which the bacterium protects the beetle eggs against pathogenic fungi by producing different bioactive secondary metabolites [77, 78]. The versatile transmission, infection, and symbiotic patterns of *B*. *gladioli* raise the possibility that it might have been introduced into Bangladesh not only through rice imports but also through imports of other agricultural products or through other human-mediated routes (e.g., international travellers carrying USA-linked strains). To determine the exact routes of introduction of pathogenic *B. gladioli* strains into Bangladesh, comparative genomic analyses including *B*. *gladioli* strains from neighboring countries—e.g. India, Myanmar, China, Pakistan and Thailand—will be essential. However, genomic data from rice or any other samples from these countries are currently lacking, which prevents us from inferring potential introductions from these countries. If these regions harbor genetically similar strains to the ones isolated in Bangladesh, the finding would strengthen the hypothesis that *B*. *gladioli* was introduced into Bangladesh via rice imports. Conversely, other mechanisms of human-mediated transmission may be the dominant route if the closest relatives remain linked to clinical isolates from the USA.

The 37 genes analyzed in this study are directly associated with plant pathogenicity (Table 2). Since (1) the pathogenicity mechanisms of *B. gladioli* are poorly understood, (2) *B*. *gladioli* and *B*. *glumae* are closely related to each other, and (3) previous studies have shown that most of the pathogenicity factors that we studied are highly conserved between both species [36–38], we used the most studied *B*. *glumae* strain, BGR1, as the reference for our study. Both *B*. *glumae* and *B. gladioli* cause panicle infection, supporting the hypothesis that they migrate from the seeds to vegetative tissues and ultimately to the panicles [79]. For migration, *B*. *glumae* uses flagella/motility and lipopolysaccharide-related gene clusters [60, 80]. After reaching the specific plant tissues, such as developing panicles, *B*. *glumae* deploys its main virulence factor, toxoflavin, which generates hydrogen peroxide and causes tissue damage [81, 82]. Previous studies have confirmed that most of the *B*. *gladioli* strains also produce toxoflavin and exhibit *tox* operons that are very similar to that of *B*. *glumae* BGR1 [17, 23, 31, 36, 38, 83]. Other important pathogenicity factors of *B*. *glumae* are lipases LipA and LipB, which act as cell wall-degrading enzymes [84]. The presence of a *lipA* ortholog has been confirmed in 7 *B*. *gladioli* strains [36, 37], and we identified orthologs of both *lipA* and *lipB* in nearly all *B. gladioli* strains (Table S3). Endopolygalacturonases PehA and PehB, secreted through the type II secretion system (T2SS), may not be directly involved in the initial stages of the disease caused by *B*. *glumae* but may have a role at a later stage of the infection process [85]. In a study of four *B. gladioli* strains, *pehB* was found in all four strains, whereas *pehA* was found in only one [37]; in contrast, we found *pehB* and *pehA* in 8% and 72% of the strains included in our study, respectively (Table S3). The TofI/TofR quorum-sensing system, mediated by the C8-HSL signal, regulates toxoflavin biosynthesis, lipase production, flagellar biosynthesis, and motility in *B*. *glumae* [60, 84, 85]. The TofI/TofR quorum-sensing system has also been identified in *B. gladioli* [38]. Our analyses indicate that almost every *B*. *gladioli* strain possessed quorum-sensing, toxoflavin, flagella, and lipase synthesis genes (Table S3), suggesting that the mechanisms used by *B*. *gladioli* to cause BPB are largely similar to those used by *B. glumae*.

Most of the *B. gladioli* strains included in our study exhibit similar virulence-related gene contents (Table S3), which may explain why the pathogen can switch hosts relatively easily. In fact, in the case of *Pseudomonas aeruginosa* PA14—a human pathogen associated with cystic fibrosis that is closely related to *Burkholderia* and that is also capable of infecting plants—it has been reported that many of the virulence factors that are essential for animal infection also contribute to plant pathogenicity [86]. Nonetheless, we found some genetic differences between human- and plant-associated *B. gladioli* strains. In particular, our Monte Carlo tests were significant for two of the studied genes (*tsr* and *motB*, *P* = 0.0011), indicating that human-to-plant host transitions frequently co-occurred with losses of these two genes (Table S3). That suggests that *tsr* and *motB* are more important for human pathogenicity than for plant pathogenicity, and that loss of these genes was often adaptive after human-to-plant transitions.

Both *tsr* and *motB* are important for bacterial motility. Tsr is a transmembrane receptor protein that detects serine as an attractant. It has a periplasmic ligand-binding domain that monitors attractant levels in the environment and a cytoplasmic signaling domain that alters the rotational bias (clockwise/anticlockwise) of flagellar motors [87–89]. The absence of the *tsr* gene in 26% of the studied plant-derived strains may result in a defective chemotactic response and inhibition of motility in these strains. Indeed, a mutation in the signaling domain of Tsr inhibiting the CheA kinase abolished swarming motility in *Escherichia coli* [90]. In addition, in *E. coli*, Tsr mediates adhesion, colonization, and transmigration in the lung mucosal barrier [91] and facilitates Enterobacteriaceae to migrate toward enterohemorrhagic lesions to use human serum as a source of nutrients [92]. In *Salmonella typhimurium*, *tsr* mutants grow less efficiently than wild-type strains, yet a bioinformatic analysis of 100,000 genomes identified *tsr* as one of the most frequently mutated loci [93], suggesting that certain unidentified environments favor these mutants [94]. In terms of plant-pathogenic *Burkholderia*, the mutation of *tsr* in the *B*. *glumae* BGR1 strain did not affect swarming motility, but it did impair swimming motility in the mutants. Additionally, in the *B*. *glumae* 336gr-1 strain, *tsr* was found to exhibit a 78 bp deletion and reported dispensable for plant pathogenicity [61]. Therefore, the absence of the *tsr* gene in some of the plant-pathogenic *B*. *gladioli* strains may be a strategy for adaptation through the loss of dispensable genetic elements.

The MotA and MotB proteins form the stator (the nonrotating part) of the flagellar motor anchored around the basal body [95–97]. The two proteins combine to create an ion channel that allows proton or ion flow, inducing a conformational change in MotA. This conformational change allows MotA to interact with FliG, generating torque to drive flagellar rotation [98–101]. Since 23% of the plant-derived *B*. *gladioli* strains lack the *motB* gene, those strains may show a lack of flagellar rotation and/or impaired motility. Indeed, deletion of the *motB* gene in *E*. *coli* results in loss of flagellar rotation [102]. Recently, there has been an increase in the discovery of mutants with a *mot* phenotype [103–106], which were previously rare [107]. In addition, in *Salmonella*, a mutation affecting the highly conserved Asp33 residue of MotB resulted in large speed fluctuations and frequent pausing of motor rotation at low load [108]. Recent findings in *Burkholderia cenocepacia* further confirmed that MotB has a similar role in the genus *Burkholderia* by showing that ΔmotB mutants lose flagellar movement and complementation restores motility [109]. This finding suggests that MotB probably plays a similar role in *B*. *gladioli*.

Loss of the *tsr* and/or *motB* genes from some of the plant pathogenic *B*. *gladioli* strains may have resulted in reduced or no chemotactic response and/or motility in these strains. Impaired or reduced motility can lead to the induction of surface adhesins and increased biofilm formation [104]. Indeed, some motility (*mot*) defective *Bacillus* strains exhibit both flagellar impairment and increased biofilm formation [104, 107]. In many plant-pathogenic bacteria, biofilm formation has evolved as an adaptive strategy to successfully achieve host colonization [110]. Many species of the *Burkholderia* genus are known to produce biofilms; for instance, plant and human pathogenic strains of *B. glumae* have been reported to produce biofilms in NB medium [111], and plant pathogenic strains of *B. plantarii* and *B. gladioli* have been shown to produce biofilms in PDB medium [112, 113]. These observations lend support to the possibility that loss of *tsr* and/or *motB* in some plant pathogenic *B. gladioli* strains may have resulted in an adaptive increase in biofilm formation. Biofilms stabilize the colonization on the plant and protect bacteria from different host and environmental stresses such as UV radiation, osmotic stresses, desiccation, and components of the host immune system [114–116]. Moreover, biofilms facilitate horizontal gene transfer among bacteria living therein [117, 118], aiding in the acquisition of antibiotic resistance [119]. However, bacterial biofilms facilitate survival not only on plants, but also on other environments, including surfaces and human tissues [120]. Indeed, Bcc clinical isolates tend to exhibit stronger biofilm formation than Bcc agricultural isolates [121], and the human-derived *B. glumae* strain AU6208 showed increased biofilm formation compared with the rice pathogenic *B*. *glumae* LMG 2196 strain [111]. Thus, it remains unclear why losses of *tsr* and *motB* tend to co-occur with human-to-plant transitions specifically.

## CONCLUSION

In conclusion, our phylogenetic analysis suggests five independent introduction events of pathogenic *B. gladioli* strains into Bangladesh, likely occurring through international trade or other human-mediated mechanisms. Four of these introductions appear to have occurred from the USA, either directly or indirectly. No evidence of inter-district dissemination was found. The co-occurrence of *tsr* and *motB* gene losses with human-to-plant host shifts suggests that this may represent an adaptive strategy for plant-pathogenic *B. gladioli*. Our findings may help improve the understanding of pathogenicity mechanisms and support the development of sustainable management strategies. In particular, our findings may guide agricultural quarantine policies to limit cross-border transmission and inform clinical monitoring to identify genetic markers associated with host adaptation and virulence, thereby aiding in the management of *B. gladioli* in both agricultural and clinical settings.

## LIST OF ABBREVIATIONS

*ANI*: Average Nucleotide Identity
*BPB*: Bacterial Panicle Blight
*Bcc*: Burkholderia cepacia complex
*CNV*: Copy number variation
*FET*: Fisher’s Exact Test
*GTR*: Generalized Time Reversible
*iTOL*: Interactive Tree of Life
*KBA*: King’s B agar
*PDB*: Potato Dextrose Broth
*T2SS*: type II secretion system
*T3SS*: type III secretion system
*T6SS*: type VI secretion system

## DECLARATIONS

### Ethics approval and consent to participate

Site access permissions were obtained in advance from the landowners of the surveyed fields. Permission was also obtained to ship DNA samples from the Department of Plant Pathology, Bangladesh Agricultural University (BAU), to the Nevada Genomics Center at the University of Nevada, Reno.

### Consent for publication

Not applicable.

### Availability of data and materials

All sequence data of the 19 strains of *B*. *gladioli* were deposited in the GenBank database under the BioProject accession number PRJNA1252168. The accession numbers of all 19 strains are provided in Table S4.

### Authors’ contributions

IAP participated in securing funding and in the conception and design of the study, participated in molecular identification, performed statistical and genomic analyses, generated visualizations and drafted the manuscript. MNU participated in securing funding and performed field isolate collection and DNA extraction. AG performed whole genome sequencing and assisted in data analysis. MRI participated in securing funding and designed and supervised field isolate collection and molecular identification. DAP participated in securing funding and in the conception and design of the study, supervised statistical and genomic analyses, and drafted the manuscript. All authors have read and approved the final version of the manuscript.

## Acknowledgements

The authors are grateful to the local personnel of the Department of Agricultural Extension, Ministry of Agriculture, Government of the People’s Republic of Bangladesh for their support during sample collection from fields.

## References

1. Depoorter E, Bull MJ, Peeters C, Coenye T, Vandamme P, Mahenthiralingam E. *Burkholderia*: an update on taxonomy and biotechnological potential as antibiotic producers. Appl Microbiol Biotechnol. 2016;100:5215–29. 10.1007/s00253-016-7520-x.

2. Carson LA, Favero MS, Bond WW, Petersen NJ. Morphological, Biochemical, and Growth Characteristics of *Pseudomonas cepacia* from Distilled Water. Appl Microbiol. 1973;25:476–83. 10.1128/am.25.3.476-483.1973.

3. Bevivino A, Tabacchioni S, Chiarini L, Carusi MV, Del Gallo M, Visca P. Phenotypic comparison between rhizosphere and clinical isolates of *Burkholderia cepacia*. Microbiology. 1994;140:1069–77. 10.1099/13500872-140-5-1069.

4. Homma Y, Chikuo Y, Ogoshi A. Mode of suppression of sugar beet damping-off caused by Rhizoctonia solani by seed bacterization with *Pseudomonas cepacia*. Bull SROP. 1991;14:115–8.

5. Jones C, Webster G, Mullins AJ, Jenner M, Bull MJ, Dashti Y, et al. Kill and cure: genomic phylogeny and bioactivity of *Burkholderia gladioli* bacteria capable of pathogenic and beneficial lifestyles. Microb Genomics. 2021;7. 10.1099/mgen.0.000515.

6. Vandamme P, Govan JR, LiPuma JJ. Burkholderia : molecular microbiology and genomics. Wymondham, UK: Horizon Bioscience; 2007.

7. Estrada-de Los Santos P, Vinuesa P, Martínez-Aguilar L, Hirsch AM, Caballero-Mellado J. Phylogenetic Analysis of *Burkholderia* Species by Multilocus Sequence Analysis. Curr Microbiol. 2013;67:51–60. 10.1007/s00284-013-0330-9.

8. Mahenthiralingam E, Urban TA, Goldberg JB. The multifarious, multireplicon *Burkholderia cepacia* complex. Nat Rev Microbiol. 2005;3:144–56. 10.1038/nrmicro1085.

9. Lee H-H, Park J, Jung H, Seo Y-S. Pan-Genome Analysis Reveals Host-Specific Functional Divergences in *Burkholderia gladioli*. Microorganisms. 2021;9:1123. 10.3390/microorganisms9061123.

10. Palieroni NJ. Genus I. Pseudomonas Migula 1894, 237AL (nom. cons. Opin. 5, Jud. Comm. 1952, 237). In: Bergey’s Manual of Systematic Bacteriology. Baltimore, MD: Williams & Wilkins; 1984. p. 141–99.

11. Wright PJ, Clark RG, Hale CN. A storage soft rot of New Zealand onions caused by *Pseudomonas gladioli* pv. *alliicola*. N Z J Crop Hortic Sci. 1993;21:225–7. 10.1080/01140671.1993.9513773.

12. Hildebrand DC, Palleroni NJ, Doudoroff M. Synonymy of *Pseudomonas gladioli* Severini 1913 and Pseudomonas marginata (McCulloch 1921) Stapp 1928. Int J Syst Bacteriol. 1973;23:433–7. 10.1099/00207713-23-4-433.

13. Gill WM, Tsuneda A. The interaction of the soft rot bacterium *Pseudomonas gladioli* pv. *agaricicola* with Japanese cultivated mushrooms. Can J Microbiol. 1997;43:639–48. 10.1139/m97-091.

14. Van Damme PA, Johannes AG, Cox HC, Berends W. On toxoflavin, the yellow poison of *pseudomonas cocovenenans*. Recl Trav Chim Pays-Bas. 1960;79:255–67. 10.1002/recl.19600790305.

15. Zhou F, Ning H, Chen F, Wu W, Chen A, Zhang J. *Burkholderia gladioli* infection isolated from the blood cultures of newborns in the neonatal intensive care unit. Eur J Clin Microbiol Infect Dis. 2015;34:1533–7. 10.1007/s10096-015-2382-1.

16. Zanotti C, Munari S, Brescia G, Barion U. *Burkholderia gladioli* sinonasal infection. Eur Ann Otorhinolaryngol Head Neck Dis. 2019;136:55–6. 10.1016/j.anorl.2018.01.011.

17. Nandakumar R, Shahjahan AKM, Yuan XL, Dickstein ER, Groth DE, Clark CA, et al. *Burkholderia glumae* and *B. gladioli* Cause Bacterial Panicle Blight in Rice in the Southern United States. Plant Dis. 2009;93:896–905. 10.1094/PDIS-93-9-0896.

18. Ura H. Properties of the bacteria isolated from rice plants with leaf-sheath browning and grain rot. Ann Phytopathol Soc Jpn. 1996;62:640.

19. Cottyn B, Cerez MT, van Outryve MF, Barroga J, Swings J, Mew TW. Bacterial Diseases of Rice. I. Pathogenic Bacteria Associated with Sheath Rot Complex and Grain Discoloration of Rice in the Philippines. Plant Dis. 1996;80:429. 10.1094/PD-80-0429.

20. Yuan X. Identification of Bacterial Pathogens Causing Panicle Blight of Rice in Louisiana. La State Univ Agric Mech Coll. 2004.

21. Mulaw T, Wamishe Y, Jia Y. Characterization and in Plant Detection of Bacteria That Cause Bacterial Panicle Blight of Rice. Am J Plant Sci. 2018;09:667–84. 10.4236/ajps.2018.94053.

22. Lu SE, Allen TW. Causal Agents of Bacterial Panicle Blight of Rice and Evaluation of Disease Resistance of Rice Cultivars to the Disease in Mississippi. 34th Proc Rice Tech Work Group RTWG. 2012;:76.

23. Nandakumar R, Rush MC, Correa F. Association of *Burkholderia glumae* and *B. gladioli* with Panicle Blight Symptoms on Rice in Panama. Plant Dis. 2007;91:767–767. 10.1094/PDIS-91-6-0767C.

24. Riera-Ruiz C, Vargas J, Cedeño C, Quirola P, Escobar M, Cevallos-Cevallos JM, et al. First Report of *Burkholderia glumae* Causing Bacterial Panicle Blight on Rice in Ecuador. Plant Dis. 2014;98:988–988. 10.1094/PDIS-10-13-1024-PDN.

25. Mirghasempour SA, Huang S, Xie GL. First Report of *Burkholderia gladioli* Causing Rice Panicle Blight and Grain Discoloration in China. Plant Dis. 2018;102:2635. 10.1094/PDIS-05-18-0758-PDN.

26. Md-Zali AZ, Ja’afar Y, Paramisparan K, Ismail SI, Saad N, Hata EM, et al. First Report of *Burkholderia gladioli* Causing Bacterial Panicle Blight of Rice in Malaysia. Plant Dis. 2023;107:551. 10.1094/PDIS-03-22-0650-PDN.

27. Goto K. A new bacterial disease of rice. Ann Phytopathol Soc Jpn. 1956;21:46–7.

28. Azegami K, Nishiyama K, Watanabe Y, Kadota I, Ohuchi A, Fukazawa C. *Pseudomonas plantarii* sp. nov., the Causal Agent of Rice Seedling Blight. Int J Syst Bacteriol. 1987;37:144– 52. 10.1099/00207713-37-2-144.

29. Tsushima S. Epidemiology of bacterial grain rot of rice caused by *Pseudomonas glumae*. JARQ. 1996;30:85–9.

30. Zhou X-G. Sustainable Strategies for Managing Bacterial Panicle Blight. In: Protecting Rice Grains in the Post-Genomic Era. 2019. p. 67.

31. Islam MdR, Jannat R, Protic IA, Happy MstNA, Samin SI, Mita MM, et al. First Report of Bacterial Panicle Blight in Rice Caused by *Burkholderia gladioli* in Bangladesh. Plant Dis. 2023;107:2837. 10.1094/PDIS-02-23-0229-PDN.

32. Saha I, Durand-Morat A, Nalley LL, Alam MJ, Nayga R. Rice quality and its impacts on food security and sustainability in Bangladesh. PLOS ONE. 2021;16:e0261118. 10.1371/journal.pone.0261118.

33. Protic IA, Uddin MdN, Tushar ASMd, Auyon ST, Alvarez-Ponce D, Islam MdR. First complete genome sequence of a Bacterial Panicle Blight causing pathogen, *Burkholderia glumae*, isolated from symptomatic rice grains from Bangladesh. BMC Genomic Data. 2024;25:73. 10.1186/s12863-024-01255-5.

34. Uddin MdN, Protic IA, Md. Tushar AS, Hasan M, Saha P, Singha UR, et al. First Report of *Burkholderia glumae* Causing Bacterial Panicle Blight in Rice in Bangladesh. Plant Dis. 2025;109:491. 10.1094/PDIS-04-24-0904-PDN.

35. Mondal KK, Mani C, Verma G. Emergence of Bacterial Panicle Blight Caused by *Burkholderia glumae* in North India. Plant Dis. 2015;99:1268–1268. 10.1094/PDIS-01-15-0094-PDN.

36. Fory PA, Triplett L, Ballen C, Abello JF, Duitama J, Aricapa MG, et al. Comparative Analysis of Two Emerging Rice Seed Bacterial Pathogens. Phytopathology®. 2014;104:436–44. 10.1094/PHYTO-07-13-0186-R.

37. Seo Y-S, Lim JY, Park J, Kim S, Lee H-H, Cheong H, et al. Comparative genome analysis of rice-pathogenic *Burkholderia* provides insight into capacity to adapt to different environments and hosts. BMC Genomics. 2015;16:349. 10.1186/s12864-015-1558-5.

38. Lee J, Park J, Kim S, Park I, Seo Y. Differential regulation of toxoflavin production and its role in the enhanced virulence of *Burkholderia gladioli*. Mol Plant Pathol. 2016;17:65–76. 10.1111/mpp.12262.

39. King EO, Ward MK, Raney DE. Two simple media for the demonstration of pyocyanin and fluorescin. J Lab Clin Med. 1954;44:301–7.

40. Maeda Y, Shinohara H, Kiba A, Ohnishi K, Furuya N, Kawamura Y, et al. Phylogenetic study and multiplex PCR-based detection of *Burkholderia plantarii*, *Burkholderia glumae* and *Burkholderia gladioli* using *gyrB* and *rpoD* sequences. Int J Syst Evol Microbiol. 2006;56:1031– 8. 10.1099/ijs.0.64184-0.

41. Libuit KG, Doughty EL, Otieno JR, Ambrosio F, Kapsak CJ, Smith EA, et al. Accelerating bioinformatics implementation in public health. Microb Genomics. 2023;9. 10.1099/mgen.0.001051.

42. Petit RA, Read TD. Bactopia: a Flexible Pipeline for Complete Analysis of Bacterial Genomes. mSystems. 2020;5:e00190–20. 10.1128/mSystems.00190-20.

43. Bolger AM, Lohse M, Usadel B. Trimmomatic: a flexible trimmer for Illumina sequence data. Bioinformatics. 2014;30:2114–20. 10.1093/bioinformatics/btu170.

44. Seeman T. Shovill—Assemble Bacterial Isolate Genomes from Illumina Paired-End Reads. 2020. https://github.com/tseemann/shovill.

45. Lumpe J, Gumbleton L, Gorzalski A, Libuit K, Varghese V, Lloyd T, et al. GAMBIT (Genomic Approximation Method for Bacterial Identification and Tracking): A methodology to rapidly leverage whole genome sequencing of bacterial isolates for clinical identification. PLOS ONE. 2023;18:e0277575. 10.1371/journal.pone.0277575.

46. Seemann T. Prokka: rapid prokaryotic genome annotation. Bioinformatics. 2014;30:2068–9. 10.1093/bioinformatics/btu153.

47. Waterhouse RM, Seppey M, Simão FA, Manni M, Ioannidis P, Klioutchnikov G, et al. BUSCO Applications from Quality Assessments to Gene Prediction and Phylogenomics. Mol Biol Evol. 2018;35:543–8. 10.1093/molbev/msx319.

48. Sayers EW, Cavanaugh M, Clark K, Ostell J, Pruitt KD, Karsch-Mizrachi I. GenBank. Nucleic Acids Res. 2019;47:D94–9. 10.1093/nar/gky989.

49. Jain C, Dilthey A, Koren S, Aluru S, Phillippy AM. A Fast Approximate Algorithm for Mapping Long Reads to Large Reference Databases. J Comput Biol. 2018;25:766–79. 10.1089/cmb.2018.0036.

50. Shimoyama Y. ANIclustermap: A tool for drawing ANI clustermap between all-vs-all microbial genomes [Computer software]. 2022. https://github.com/moshi4/ANIclustermap.

51. Page AJ, Cummins CA, Hunt M, Wong VK, Reuter S, Holden MTG, et al. Roary: rapid large-scale prokaryote pan genome analysis. Bioinformatics. 2015;31:3691–3. 10.1093/bioinformatics/btv421.

52. Price MN, Dehal PS, Arkin AP. FastTree 2 – Approximately Maximum-Likelihood Trees for Large Alignments. PLoS ONE. 2010;5:e9490. 10.1371/journal.pone.0009490.

53. Letunic I, Bork P. Interactive Tree of Life (iTOL) v6: recent updates to the phylogenetic tree display and annotation tool. Nucleic Acids Res. 2024;52:W78–82. 10.1093/nar/gkae268.

54. O’Leary NA, Wright MW, Brister JR, Ciufo S, Haddad D, McVeigh R, et al. Reference sequence (RefSeq) database at NCBI: current status, taxonomic expansion, and functional annotation. Nucleic Acids Res. 2016;44:D733–45. 10.1093/nar/gkv1189.

55. Fischer S, Brunk BP, Chen F, Gao X, Harb OS, Iodice JB, et al. Using OrthoMCL to Assign Proteins to OrthoMCL-DB Groups or to Cluster Proteomes Into New Ortholog Groups. Curr Protoc Bioinforma. 2011;35. 10.1002/0471250953.bi0612s35.

56. Ham JH, Melanson RA, Rush MC. *Burkholderia glumae* : next major pathogen of rice? Mol Plant Pathol. 2011;12:329–39. 10.1111/j.1364-3703.2010.00676.x.

57. Kim J, Oh J, Choi O, Kang Y, Kim H, Goo E, et al. Biochemical Evidence for ToxR and ToxJ Binding to the *tox* Operons of *Burkholderia glumae* and Mutational Analysis of ToxR. J Bacteriol. 2009;191:4870–8. 10.1128/JB.01561-08.

58. Knapp A, Voget S, Gao R, Zaburannyi N, Krysciak D, Breuer M, et al. Mutations improving production and secretion of extracellular lipase by *Burkholderia glumae* PG1. Appl Microbiol Biotechnol. 2016;100:1265–73. 10.1007/s00253-015-7041-z.

59. De Paula Lelis T. Characterization of the Integrated Signaling Network of *Burkholderia Glumae* for the Regulation of Virulence-Related function in the Bacterial Pathogenesis of Rice Plants. Doctor of Philosophy. Louisiana State University and Agricultural and Mechanical College; 2019. 10.31390/gradschool_dissertations.4996.

60. Kim J, Kang Y, Choi O, Jeong Y, Jeong J, Lim JY, et al. Regulation of polar flagellum genes is mediated by quorum sensing and FlhDC in *Burkholderia glumae*. Mol Microbiol. 2007;64:165–79. 10.1111/j.1365-2958.2007.05646.x.

61. Francis F, Kim J, Ramaraj T, Farmer A, Rush MC, Ham JH. Comparative genomic analysis of two *Burkholderia glumae* strains from different geographic origins reveals a high degree of plasticity in genome structure associated with genomic islands. Mol Genet Genomics. 2013;288:195–203. 10.1007/s00438-013-0744-x.

62. Jang MS, Goo E, An JH, Kim J, Hwang I. Quorum Sensing Controls Flagellar Morphogenesis in *Burkholderia glumae*. PLoS ONE. 2014;9:e84831. 10.1371/journal.pone.0084831.

63. Kim S, Park J, Choi O, Kim J, Seo Y-S. Investigation of Quorum Sensing-Dependent Gene Expression in *Burkholderia gladioli* BSR3 through RNA-seq Analyses. J Microbiol Biotechnol. 2014;24:1609–21. 10.4014/jmb.1408.08064.

64. Silva PRAD, Vidal MS, Soares CDP, Polese V, Tadra-Sfeir MZ, Souza EMD, et al. Sugarcane apoplast fluid modulates the global transcriptional profile of the diazotrophic bacteria *Paraburkholderia tropica* strain Ppe8. PLOS ONE. 2018;13:e0207863. 10.1371/journal.pone.0207863.

65. Bouteiller M, Dupont C, Bourigault Y, Latour X, Barbey C, Konto-Ghiorghi Y, et al. *Pseudomonas* Flagella: Generalities and Specificities. Int J Mol Sci. 2021;22:3337. 10.3390/ijms22073337.

66. Benjamini Y, Hochberg Y. Controlling the False Discovery Rate: A Practical and Powerful Approach to Multiple Testing. J R Stat Soc Ser B Stat Methodol. 1995;57:289–300. 10.1111/j.2517-6161.1995.tb02031.x.

67. Maddison WP, FitzJohn RG. The Unsolved Challenge to Phylogenetic Correlation Tests for Categorical Characters. Syst Biol. 2015;64:127–36. 10.1093/sysbio/syu070.

68. Ahmed T. Bangladesh: Grain and feed update. United States Department of Agriculture; 2023.

69. Simoes AJG, Hidalgo CA. The Economic Complexity Observatory: An Analytical Tool for Understanding the Dynamics of Economic Development. In: Scalable Integration of Analytics and Visualization: Papers from the 2011 AAAI Workshop (WS-11-17). AAAI Press; 2011.

70. Govan JR, Brown AR, Jones AM. Evolving Epidemiology of *Pseudomonas Aeruginosa* and the *Burkholderia Cepacia* Complex in Cystic Fibrosis Lung Infection. Future Microbiol. 2007;2:153–64. 10.2217/17460913.2.2.153.

71. LiPuma JJ, Mortensen JE, Dasen SE, Edlind TD, Schidlow DV, Burns JL, et al. Ribotype analysis of *Pseudomonas cepacia* from cystic fibrosis treatment centers. J Pediatr. 1988;113:859–62. 10.1016/S0022-3476(88)80018-0.

72. LiPuma JJ. The Changing Microbial Epidemiology in Cystic Fibrosis. Clin Microbiol Rev. 2010;23:299–323. 10.1128/CMR.00068-09.

73. Suárez-Moreno ZR, Caballero-Mellado J, Coutinho BG, Mendonça-Previato L, James EK, Venturi V. Common Features of Environmental and Potentially Beneficial Plant-Associated *Burkholderia*. Microb Ecol. 2012;63:249–66. 10.1007/s00248-011-9929-1.

74. Peterson AE, Chitnis AS, Xiang N, Scaletta JM, Geist R, Schwartz J, et al. Clonally related *Burkholderia contaminans* among ventilated patients without cystic fibrosis. Am J Infect Control. 2013;41:1298–300. 10.1016/j.ajic.2013.05.015.

75. Lucero CA, Cohen AL, Trevino I, Rupp AH, Harris M, Forkan-Kelly S, et al. Outbreak of *Burkholderia cepacia* complex among ventilated pediatric patients linked to hospital sinks. Am J Infect Control. 2011;39:775–8. 10.1016/j.ajic.2010.12.005.

76. Hamidizade M, Taghavi SM, Soleimani A, Bouazar M, Abachi H, Portier P, et al. Wild mushrooms as potential reservoirs of plant pathogenic bacteria: a case study on *Burkholderia gladioli*. Microbiol Spectr. 2024;12:e03395–23. 10.1128/spectrum.03395-23.

77. Flórez LV, Scherlach K, Gaube P, Ross C, Sitte E, Hermes C, et al. Antibiotic-producing symbionts dynamically transition between plant pathogenicity and insect-defensive mutualism. Nat Commun. 2017;8:15172. 10.1038/ncomms15172.

78. Kaltenpoth M, Flórez LV. Versatile and Dynamic Symbioses Between Insects and *Burkholderia* Bacteria. Annu Rev Entomol. 2020;65:145–70. 10.1146/annurev-ento-011019-025025.

79. Ortega L, Rojas CM. Bacterial Panicle Blight and *Burkholderia glumae* : From Pathogen Biology to Disease Control. Phytopathology®. 2021;111:772–8. 10.1094/PHYTO-09-20-0401-RVW.

80. Lee C, Mannaa M, Kim N, Kim J, Choi Y, Kim SH, et al. Stress Tolerance and Virulence-Related Roles of Lipopolysaccharide in *Burkholderia glumae*. Plant Pathol J. 2019;35:445–58. 10.5423/PPJ.OA.04.2019.0124.

81. Latuasan HE, Berends W. On the origin of the toxicity of toxoflavin. Biochim Biophys Acta. 1961;52:502–8. 10.1016/0006-3002(61)90408-5.

82. Iiyama K, Furuya N, Takanami Y, Matsuyama N. A Role of Phytotoxin in Virulence of *Pseudomonas glumae* Kurita et Tabei. Jpn J Phytopathol. 1995;61:470–6. 10.3186/jjphytopath.61.470.

83. Nandakumar R, Bollich PA, Shahjahan AKM, Groth DE, Rush MC. Evidence for the soilborne nature of the rice sheath rot and panicle blight pathogen, Burkholderia gladioli_1_. Can J Plant Pathol. 2008;30:148–54. 10.1080/07060660809507505.

84. Devescovi G, Bigirimana J, Degrassi G, Cabrio L, LiPuma JJ, Kim J, et al. Involvement of a Quorum-Sensing-Regulated Lipase Secreted by a Clinical Isolate of *Burkholderia glumae* in Severe Disease Symptoms in Rice. Appl Environ Microbiol. 2007;73:4950–8. 10.1128/AEM.00105-07.

85. Degrassi G, Devescovi G, Kim J, Hwang I, Venturi V. Identification, characterization and regulation of two secreted polygalacturonases of the emerging rice pathogen *Burkholderia glumae*: Two secreted polygalacturonases of *Burkholderia glumae*. FEMS Microbiol Ecol. 2008;65:251–62. 10.1111/j.1574-6941.2008.00516.x.

86. Rahme LG, Stevens EJ, Wolfort SF, Shao J, Tompkins RG, Ausubel FM. Common Virulence Factors for Bacterial Pathogenicity in Plants and Animals. Science. 1995;268:1899–902. 10.1126/science.7604262.

87. Ames P, Parkinson JS. Constitutively signaling fragments of Tsr, the *Escherichia coli* serine chemoreceptor. J Bacteriol. 1994;176:6340–8. 10.1128/jb.176.20.6340-6348.1994.

88. Krikos A, Conley MP, Boyd A, Berg HC, Simon MI. Chimeric chemosensory transducers of *Escherichia coli*. Proc Natl Acad Sci. 1985;82:1326–30. 10.1073/pnas.82.5.1326.

89. Manoil C, Beckwith J. A Genetic Approach to Analyzing Membrane Protein Topology. Science. 1986;233:1403–8. 10.1126/science.3529391.

90. Burkart M, Toguchi A, Harshey RM. The chemotaxis system, but not chemotaxis, is essential for swarming motility in *Escherichia coli*. Proc Natl Acad Sci. 1998;95:2568–73. 10.1073/pnas.95.5.2568.

91. Han B, Li M, Xu Y, Islam D, Khang J, Del Sorbo L, et al. Tsr Chemoreceptor Interacts With IL-8 Provoking *E*. *coli* Transmigration Across Human Lung Epithelial Cells. Sci Rep. 2016;6:31087. 10.1038/srep31087.

92. Glenn SJ, Gentry-Lear Z, Shavlik M, Harms MJ, Asaki TJ, Baylink A. Bacterial vampirism mediated through taxis to serum. eLife. 2024;12:RP93178. 10.7554/eLife.93178.3.

93. Cherry JL. Selection-Driven Gene Inactivation in *Salmonella*. Genome Biol Evol. 2020;12:18–34. 10.1093/gbe/evaa010.

94. Gül E, Huuskonen J, Abi Younes A, Maurer L, Enz U, Zimmermann J, et al. *Salmonella* T3SS-2 virulence enhances gut-luminal colonization by enabling chemotaxis-dependent exploitation of intestinal inflammation. Cell Rep. 2024;43:113925. 10.1016/j.celrep.2024.113925.

95. Blair DF. Flagellar movement driven by proton translocation. FEBS Lett. 2003;545:86–95. 10.1016/S0014-5793(03)00397-1.

96. Weiss LE, Badalamenti JP, Weaver LJ, Tascone AR, Weiss PS, Richard TL, et al. Engineering motility as a phenotypic response to LuxI/R-dependent quorum sensing in *Escherichia coli*. Biotechnol Bioeng. 2008;100:1251–5. 10.1002/bit.21862.

97. Asai Y, Kojima S, Kato H, Nishioka N, Kawagishi I, Homma M. Putative channel components for the fast-rotating sodium-driven flagellar motor of a marine bacterium. J Bacteriol. 1997;179:5104–10. 10.1128/jb.179.16.5104-5110.1997.

98. Garza AG, Harris-Haller LW, Stoebner RA, Manson MD. Motility protein interactions in the bacterial flagellar motor. Proc Natl Acad Sci. 1995;92:1970–4. 10.1073/pnas.92.6.1970.

99. Kojima S, Blair DF. Conformational Change in the Stator of the Bacterial Flagellar Motor. Biochemistry. 2001;40:13041–50. 10.1021/bi011263o.

100. Berg HC. The Rotary Motor of Bacterial Flagella. Annu Rev Biochem. 2003;72:19–54. 10.1146/annurev.biochem.72.121801.161737.

101. Gabel CV, Berg HC. The speed of the flagellar rotary motor of *Escherichia coli* varies linearly with protonmotive force. Proc Natl Acad Sci. 2003;100:8748–51. 10.1073/pnas.1533395100.

102. Blair DF, Berg HC. Restoration of Torque in Defective Flagellar Motors. Science. 1988;242:1678–81. 10.1126/science.2849208.

103. Ko M, Park C. Two novel flagellar components and H-NS are involved in the motor function of *Escherichia coli*. J Mol Biol. 2000;303:371–82. 10.1006/jmbi.2000.4147.

104. Blair KM, Turner L, Winkelman JT, Berg HC, Kearns DB. A Molecular Clutch Disables Flagella in the *Bacillus subtilis* Biofilm. Science. 2008;320:1636–8. 10.1126/science.1157877.

105. Pilizota T, Brown MT, Leake MC, Branch RW, Berry RM, Armitage JP. A molecular brake, not a clutch, stops the *Rhodobacter sphaeroides* flagellar motor. Proc Natl Acad Sci. 2009;106:11582–7. 10.1073/pnas.0813164106.

106. Boehm A, Kaiser M, Li H, Spangler C, Kasper CA, Ackermann M, et al. Second Messenger-Mediated Adjustment of Bacterial Swimming Velocity. Cell. 2010;141:107–16. 10.1016/j.cell.2010.01.018.

107. Guttenplan SB, Kearns DB. Regulation of flagellar motility during biofilm formation. FEMS Microbiol Rev. 2013;37:849–71. 10.1111/1574-6976.12018.

108. Che Y, Nakamura S, Morimoto YV, Kami-ike N, Namba K, Minamino T. Load-sensitive coupling of proton translocation and torque generation in the bacterial flagellar motor. Mol Microbiol. 2014;91:175–84. 10.1111/mmi.12453.

109. Lewis JM, Jebeli L, Coulon PML, Lay CE, Scott NE. Glycoproteomic and proteomic analysis of *Burkholderia cenocepacia* reveals glycosylation events within FliF and MotB are dispensable for motility. Microbiol Spectr. 2024;12:e00346–24. 10.1128/spectrum.00346-24.

110. Castiblanco LF, Sundin GW. New insights on molecular regulation of biofilm formation in plant-associated bacteria. J Integr Plant Biol. 2016;58:362–72. 10.1111/jipb.12428.

111. Cui Z, Wang S, Kakar KU, Xie G, Li B, Chen G, et al. Genome Sequence and Adaptation Analysis of the Human and Rice Pathogenic Strain *Burkholderia glumae* AU6208. Pathogens. 2021;10:87. 10.3390/pathogens10020087.

112. Kang M, Lee D, Mannaa M, Han G, Choi H, Lee S, et al. Impact of Quorum Sensing on the Virulence and Survival Traits of *Burkholderia plantarii*. Plants. 2024;13:2657. 10.3390/plants13182657.

113. Jiao R, Zhang X, Wang Y, Ren Y, Ou D, Ling N, et al. Prevalence, phenotypic characteristics and genetic diversity of *Burkholderia gladioli* isolated from edible mushrooms. LWT. 2024;194:115805. 10.1016/j.lwt.2024.115805.

114. Costerton JW, Stewart PS, Greenberg EP. Bacterial Biofilms: A Common Cause of Persistent Infections. Science. 1999;284:1318–22. 10.1126/science.284.5418.1318.

115. Del Pozo JL, Patel R. The Challenge of Treating Biofilm-associated Bacterial Infections. Clin Pharmacol Ther. 2007;82:204–9. 10.1038/sj.clpt.6100247.

116. Yaron S, Römling U. Biofilm formation by enteric pathogens and its role in plant colonization and persistence. Microb Biotechnol. 2014;7:496–516. 10.1111/1751-7915.12186.

117. Abe K, Nomura N, Suzuki S. Biofilms: hot spots of horizontal gene transfer (HGT) in aquatic environments, with a focus on a new HGT mechanism. FEMS Microbiol Ecol. 2020;96:fiaa031. 10.1093/femsec/fiaa031.

118. Ma L, Konkel ME, Lu X. Antimicrobial Resistance Gene Transfer from *Campylobacter jejuni* in Mono- and Dual-Species Biofilms. Appl Environ Microbiol. 2021;87:e00659–21. 10.1128/AEM.00659-21.

119. Benveniste R, Davies J. Aminoglycoside Antibiotic-Inactivating Enzymes in Actinomycetes Similar to Those Present in Clinical Isolates of Antibiotic-Resistant Bacteria. Proc Natl Acad Sci. 1973;70:2276–80. 10.1073/pnas.70.8.2276.

120. Vishwakarma V. Impact of environmental biofilms: Industrial components and its remediation. J Basic Microbiol. 2020;60:198–206. 10.1002/jobm.201900569.

121. Ibrahim M, Tang Q, Shi Y, Almoneafy A, Fang Y, Xu L, et al. Diversity of potential pathogenicity and biofilm formation among *Burkholderia cepacia* complex water, clinical, and agricultural isolates in China. World J Microbiol Biotechnol. 2012;28:2113–23. 10.1007/s11274-012-1016-3.

